# Quantitative approaches to variant classification increase the yield and precision of genetic testing in Mendelian diseases: The case of hypertrophic cardiomyopathy

**DOI:** 10.1101/381467

**Authors:** Roddy Walsh, Francesco Mazzarotto, Nicola Whiffin, Rachel Buchan, William Midwinter, Alicja Wilk, Nicholas Li, Leanne Felkin, Nathan Ingold, Risha Govind, Mian Ahmad, Erica Mazaika, Mona Allouba, Xiaolei Zhang, Antonio de Marvao, Sharlene M Day, Euan Ashley, Steven D Colan, Michelle Michels, Alexandre C Pereira, Daniel Jacoby, Carolyn Y Ho, Kate L Thomson, Hugh Watkins, Paul JR Barton, Iacopo Olivotto, Stuart A Cook, James S Ware

## Abstract

**Background:** International guidelines for variant interpretation in Mendelian disease set stringent criteria to report a variant as (likely) pathogenic, prioritising control of false positive rate over test sensitivity and diagnostic yield. Genetic testing is also more likely informative in individuals with well-characterised variants from extensively studied European-ancestry populations. Inherited cardiomyopathies are relatively common Mendelian diseases that allow empirical calibration and assessment of this framework.

**Results:** We compared rare variants in large hypertrophic cardiomyopathy (HCM) cohorts to reference populations to identify variant classes with high prior likelihoods of pathogenicity, as defined by etiological fraction (EF). Analysis of variant distribution identified regions in which variants are significantly enriched in cases and variant location was a better discriminator of pathogenicity than generic computational functional prediction algorithms. Non-truncating variant classes with an EF≥0.95, and therefore clinically actionable, were identified in 5 established HCM genes. Applying this approach leads to an estimated 14-20% increase in cases with actionable HCM variants.

**Conclusions:** When found in a patient confirmed to have disease, novel variants in some genes and regions are empirically shown to have a sufficiently high probability of pathogenicity to support a “likely pathogenic” classification, even without additional segregation or functional data. This could increase the yield of high confidence actionable variants, consistent with the framework and recommendations of current guidelines. The techniques outlined offer a consistent, unbiased and equitable approach to variant interpretation for Mendelian disease genetic testing. We propose adaptations to ACMG/AMP guidelines to incorporate such evidence in a quantitative and transparent manner.

## BACKGROUND

Advances in sequencing technology have dramatically expanded the scope for genetic testing in rare Mendelian diseases, but have exposed variant interpretation as a key limiting factor for clinical application. In an effort to standardise variant assessment in clinical settings, guidelines from the American College of Medical Genetics and Genomics/Association for Molecular Pathology (ACMG/AMP) were produced in 2015[1] and have now been widely adopted[2]. These were in part prompted by the plethora of erroneous variant-disease associations in the research literature[3, 4] and the increasing realisation that individually rare variants are collectively common for many genes, as highlighted by population datasets such as the Exome Aggregation Consortium (ExAC)[5]. A critical objective of the guidelines is to limit false positive results in clinical genetic testing in order to avoid genetic misdiagnosis or false reassurance through predictive testing of a variant that is not causal.

The ACMG/AMP guidelines outline how different lines of evidence should be assessed when interpreting a variant, and the strength of evidence required for a pathogenic (or likely pathogenic) classification. However, they are deliberately broad in scope, with the intention that individual rules would be interpreted and adapted for specific diseases within the overall framework[6]. They are conservative in nature and require substantial evidence in order to classify a variant as disease-causing. In practice, while novel truncating variants can be classified as pathogenic (when found in a gene where loss of function is a known mechanism of disease and fulfilling other conditions such as rarity), variant-specific evidence (such as segregation in the family or prior functional evidence of pathogenicity) is required for non-truncating variants to be deemed actionable.

We have recently shown that clinical laboratories utilising these stringent approaches to variant classification are, as expected, under-calling pathogenic variants in well-established cardiomyopathy genes[3], prioritising high specificity at a cost of test sensitivity. Clinical outcome data from the SHaRe registry of hypertrophic cardiomyopathy (HCM) patients supports this finding, as patients with variants of uncertain significance (VUS) had outcomes intermediate between genotype-positive and negative patients, indicating a substantial proportion are likely to be pathogenic [in press at Circulation]. Some diseases, including cardiomyopathies, are highly genetically heterogeneous with thousands of distinct causative variants, many of which are private or only detected in a handful of families, so interpretation of previously unseen variants is essential to provide a molecular diagnosis to many patients. As a consequence, genetic testing can be somewhat of a lottery for patients, with a positive result often dependent on whether the putative causative variant has been previously identified and characterised.

Furthermore, the degree of certainty required to consider a specific variant causal in an individual depends on the use of that information. While predictive testing or pre-implantation genetic diagnosis requires a high degree of confidence, some treatment decisions may be made at lower confidence. In early onset diabetes, a potentially causative variant suggesting possible MODY (maturity onset diabetes of the young) might trigger a trial of sulfonylureas even if formally a VUS[7]. Dilated cardiomyopathy due to variation in lamin a/c is associated with a poor prognosis, with a propensity for life-threatening arrhythmia. A lower threshold for primary prevention ICD implantation may be adopted if a novel variant in *LMNA* is identified, even if formally classified as a VUS and predictive testing would not be undertaken on the same variant[8–10].

The likelihood of obtaining a definitive result is also dependent on the ethnicity of the patient. Data from the Partners Laboratory of Molecular Medicine (LMM) in the United States showed that Caucasian patients are more likely to get a positive result in cardiomyopathy genetic testing than “underrepresented minorities” (including African-Americans and Hispanics) and that the proportion of patients with inconclusive results was significantly greater in both Asians and “underrepresented minorities” compared to Caucasians [11]. Similar findings were observed specifically for HCM - the proportion of positive/uncertain results was 34.7%/13.9% for Caucasians and 24.2%/20.6% for non-Caucasians (p<0.0001) in the LMM cohort (n=2,912)[12]. One of the likely reasons for this discrepancy is that much of the research and clinical testing in this condition has been done in Caucasian-majority populations and therefore Caucasians are more likely to have a causative variant that has been previously characterised. Inequalities in healthcare provision and access to genetic testing in the US may also exacerbate this disparity[13]. While more genetic research in non-Caucasian populations is clearly required, these findings underline the need for improved variant analysis techniques that reduce the reliance on prior characterisation of individual variants and better distinguish poorly characterised variants that have a high likelihood of pathogenicity from those that are unlikely to be disease-causing.

For genes with a significant excess of rare variation in case cohorts over the general population, the etiological fraction (EF) provides a quantitative estimate of the probability that a rare variant detected in an individual with disease is causative, and is dependent on the gene, variant class and variant location within the gene/protein. Here, we apply this approach in validated HCM genes to empirically determine the probability that a novel variant found in a case is pathogenic before considering other evidence and further expand the framework to identify sub-genic regions (“hotspots”) in which variants have an increased likelihood of being actionable. This provides a more quantitative approach to variant classification, with the aim of both increasing the yield of high confidence pathogenic variants detected in these genes and enabling a more unbiased application of genetic testing. We outline a potential framework to integrate this approach with the ACMG/AMP guidelines for genes and diseases with available case series to derive these estimates, enabling such case-control data to be utilised in a more quantitative and transparent manner. While highlighting that variant interpretation is highly dependent on the context of gene and disease, this approach is widely applicable for other Mendelian diseases for which sufficient cases have been genetically characterised.

## RESULTS

### In established HCM-associated genes, the majority of rare variants found in cases are pathogenic

We compared the prevalence of rare variants of different classes in established HCM-associated genes between HCM cases and population controls, and calculated the odds ratio (OR) for disease. From this we derived the etiological fraction (EF) which, under a Mendelian disease model, provides an estimate of the proportion of rare variants found in affected individuals that are disease-causing, and therefore the probability that an individual variant is pathogenic.

The etiological fraction (EF) and odds ratio (OR) for non-truncating and truncating variants in validated HCM genes[15] are shown in Table 1. Truncating variants in *MYBPC3*, causative in over 9% of HCM cases, have an EF>0.99. Truncating variants in other genes with an excess over ExAC are less prevalent (occurring in <0.2% of cases in each gene) but the probability that a variant found in a case is causal is nonetheless high (>0.84). While non-truncating variants are more prevalent in the general population, leading to a lower signal to noise ratio and reduced interpretative confidence for individual variants, the majority of such variants are causal, when found in an individual with confirmed disease. However, at the gene level, only variants in *TPM1* yield an EF≥0.95.

**Table 1:** Etiological fractions and odds ratios for established HCM genes. Displayed are rare variant frequencies (ExAC filtering allele frequency < 4x10^−5^[14]), Fisher’s exact test p-values and OR and EF values (with 95% confidence intervals) for non-truncating and truncating variants in HCM genes. The etiological fraction can be interpreted as an estimate of the probability that a rare variant, found in an individual with HCM, is causative. This suggests that the majority of variants are pathogenic, and for some genes the probability that an individual variant is pathogenic is >0.9, before considering variant-specific segregation of functional data. Only variant classes with a significant excess of variants in case cohorts over ExAC are displayed.

| Gene | Case Frequency<br>(variants/total) | ExAC Frequency<br>(variants/total) | p-value | Odds Ratio<br>(OR) | Etiological Fraction<br>(EF) |
| --- | --- | --- | --- | --- | --- |
| *Non-truncating variants* |  |  |  |  |  |
| MYH7 | 13.89% (849/6112) | 1.11% (672/60469) | <0.0001 | 14.4 (12.9-15.9) | 0.930 (0.923-0.938) |
| MYBPC3 | 9.35% (578/6179) | 1.21% (555/45794) | <0.0001 | 8.4 (7.5-9.5) | 0.881 (0.868-0.895) |
| TNNT2 | 1.69% (103/6103) | 0.15% (86/57018) | <0.0001 | 11.4 (8.5-15.2) | 0.912 (0.889-0.935) |
| TNNI3 | 2.10% (127/6047) | 0.15% (79/52607) | <0.0001 | 14.3 (10.8-18.9) | 0.930 (0.912-0.948) |
| TPM1 | 1.44% (64/4447) | 0.07% (42/58642) | <0.0001 | 20.4 (13.8-30.1) | 0.951 (0.933-0.969) |
| MYL2 | 1.03% (43/4185) | 0.11% (69/60521) | <0.0001 | 9.1 (6.2-13.3) | 0.890 (0.851-0.930) |
| MYL3 | 0.84% (35/4185) | 0.14% (85/60605) | <0.0001 | 6.0 (4.0-8.9) | 0.833 (0.772-0.895) |
| ACTC1 | 0.53% (22/4185) | 0.06% (37/60198) | <0.0001 | 8.6 (5.1-14.6) | 0.884 (0.826-0.941) |
| PLN | 0.17% (9/5440) | 0.02% (15/60475) | <0.0001 | 6.7 (2.9-15.3) | 0.850 (0.737-0.964) |
| CSRP3 | 0.62% (30/4866) | 0.19% (115/60647) | <0.0001 | 3.3 (2.2-4.9) | 0.694 (0.579-0.808) |
| FHL1 | 0.78% (16/2061) | 0.09% (53/60278) | <0.0001 | 8.9 (5.1-15.6) | 0.888 (0.826-0.949) |
| TNNC1 | 0.24% (8/3335) | 0.06% (33/59192) | 0.0013 | 4.3 (2.0-9.3) | 0.768 (0.598-0.938) |
| FLNC | 3.79% (17/448) | 2.15% (1225/56897) | 0.0314 | 1.8 (1.1-2.9) | 0.442 (0.172-0.712) |
| *Truncating variants* |  |  |  |  |  |
| MYBPC3 | 9.16% (566/6179) | 0.09% (40/45794) | <0.0001 | 115.3 (83.6-159.1) | 0.991 (0.988-0.995) |
| TNNT2 | 0.18% (11/6103) | 0.03% (17/57018) | <0.0001 | 6.1 (2.8-12.9) | 0.835 (0.722-0.948) |
| TNNI3 | 0.08% (5/6047) | 0.01% (5/52607) | 0.0019 | 8.7 (2.5-30.1) | 0.885 (0.757-1.013) |
| PLN | 0.17% (9/5440) | 0.01% (4/60475) | <0.0001 | 25.1 (7.7-81.4) | 0.960 (0.917-1.003) |
| CSRP3 | 0.14% (7/4866) | 0.02% (14/60647) | 0.0006 | 6.2 (2.5-15.5) | 0.840 (0.705-0.974) |
| FHL1 | 0.15% (3/2061) | 0.00% (0/60278) | <0.0001 | 205.0 (10.6-3969.8) | 0.995 (0.981-1.009) |

### Evaluation of missense functional prediction scores

The EF can be used to assess variant prioritisation algorithms, empirically estimating the proportion of variants that are pathogenic after applying a filter or prioritisation strategy. Some of the most commonly used tools for evaluating variants are missense functional prediction algorithms.

To initially evaluate the performance of these computational algorithms for HCM gene variants, the results of nine individual predictors (CADD, FATHMM, fMKL, LRT, mutation assessor, mutation taster, Polyphen-2, PROVEAN and SIFT) and three consensus methods (MetaLR, MetaSVM[16] and a consensus of the nine algorithms) from the dbNSFP database[17] was assessed using known pathogenic (n=298) and benign (n=349) variants in the eight sarcomeric genes (see Methods). These algorithms generally provide high sensitivity but limited specificity, as has been previously reported, although in contrast the FATHMM predictor (and MetaLR and MetaSVM consensus scores that incorporate FATHMM) have a low sensitivity for detection of pathogenic variants for *MYBPC3* and *MYL2* (Table S1). We also noted that dbNSFP does not provide predictions for certain gene/algorithm combinations (Table S2).

### Clustering analysis identifies interpretable “hot spots”, within which novel variants have a high probability of pathogenicity

For genes with an EF<0.95 for rare non-truncating variants, we examined the regional distribution of variants found in cases along the protein sequence. A novel clustering algorithm (Methods) identified a statistically significant aggregation of distinct variants (in cases) in 6 genes - *MYH7, MYBPC3, TNNI3, TNNT2, MYL3* and *CSRP3* (Figure 1, Table S3). For each cluster, the prevalence of rare variants in cases and controls was then used to calculate the EF as described above. Variants in four of these clusters *(MYH7, MYBPC3, TNNI3, TNNT2*) had an EF>0.95 (Table 2). The regions highlighted by clustering analysis correspond to key functional and protein-binding domains – the myosin motor domain of *MYH7*, troponin C and actin-binding domains in *TNNI3* and the tropomyosin-binding domain in *TNNT2. FLNC* has recently been proposed as a relatively frequent cause of HCM[18], however the high frequency of rare variation in ExAC produces only a modest EF (0.44). Although enrichment of case variants towards the C-terminus has previously been noted[18, 19], no clusters were detected in this study, though this may be due to the limited cohort size available (448 cases[19]).

**Figure 1:**
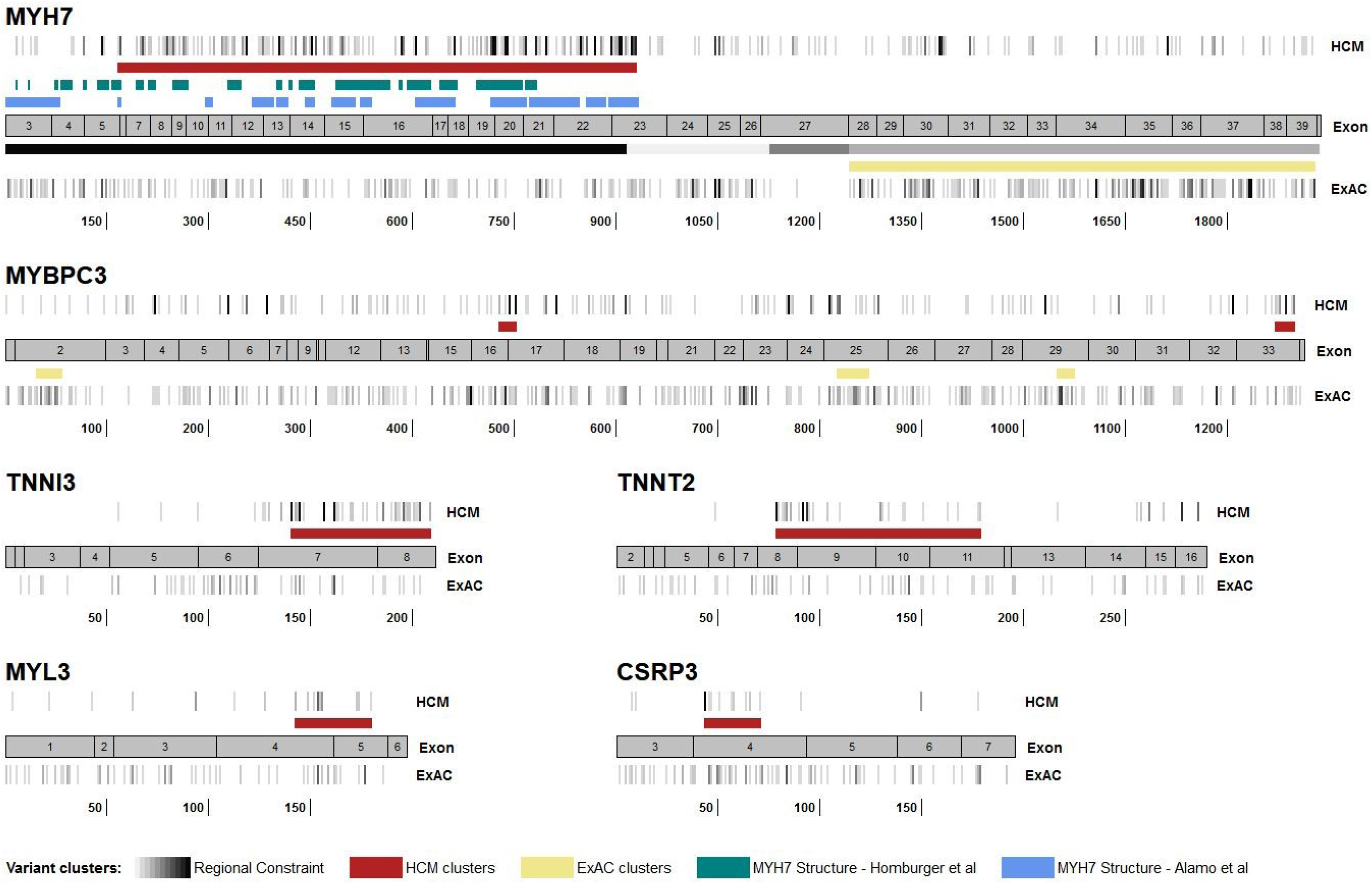
Clustering analyses identify regions enriched for disease-associated variation, and therefore within which variants have a high likelihood of pathogenicity. For six HCM genes, the location of rare missense and single amino acid inframe indel variants found in cases (all variants regardless of clinical classification) and controls are shown alongside a cartoon of the cDNA structure. Darker grey indicates higher variant density (overlapping variants not plotted separately). Regions in which variants cluster significantly in cases are shown in red, and regions with clustering in population controls (ExAC) are shown in yellow. The HCM clusters detected were: MYH7 (residues 167-931), MYBPC3 (485-502, 1248-1266), TNNI3 (141-209), TNNT2 (79-179), MYL3 (143-180) and CSRP3 (44-71). For MYH7, existing functional annotations (as described in Discussion) are superimposed: In green key residues of the converter kinetic domain and myosin mesa surface area enriched in disease-associated variants (Homburger et al[20]); in blue sites of inter- and intramolecular interaction between pairs of myosin heads (Alamo et al[21]); and in grey regions previously identified as constrained (intolerant of variation as evidenced by depletion of protein-altering variation in population controls), with the darker shades indicating higher constraint (Samocha et al[22]). The coordinates describe amino-acid position within the canonical protein sequence.

**Table 2:** Comparison of performance of variant clustering and consensus functional prediction scores in enriching for disease-associated non-truncating/missense variants in 6 HCM genes where the clustering of case variants was detected. For each gene, the EF of all rare variants is shown, followed by the EF of variants prioritised by the approach, and the EF of the remaining variants that are not prioritised. Clustering analyses identified regions of 4 genes with an EF≥0.95 (bold), and generally outperformed consensus functional prediction scores. Fisher’s exact p-values for comparison of rare variation in cases and ExAC reference samples were <0.0001 unless otherwise noted. Note – for MYBPC3 (italics), the FATHMM predictor was not included in the Consensus scores due to its poor performance for this gene, which also affected the MetaSVM and MetaLR consensus scores.

| Gene | Case Excess | EF (whole gene) | Predictor method | Prioritised variants |  | Variants not prioritised |  |
| --- | --- | --- | --- | --- | --- | --- | --- |
|  |  |  |  | Case freq. | EF | Case freq. | EF |
| MYH7 | 12.76% | 0.930 (0.923-0.938) | HCM cluster | 10.70% | 0.976 (0.972-0.981) | 3.17% | 0.746 (0.706-0.785) |
|  |  |  | Consensus | 12.55% | 0.940 (0.933-0.947) | 1.32% | 0.783 (0.728-0.839) |
|  |  |  | MetaSVM | 12.53% | 0.944 (0.937-0.951) | 1.34% | 0.739 (0.675-0.804) |
|  |  |  | MetaLR | 13.29% | 0.944 (0.938-0.951) | 0.58% (p=0.0155) | 0.406 (0.185-0.627) |
| MYBPC3 | 7.98% | 0.879 (0.865-0.893) | HCM cluster | 2.80% | 0.979 (0.971-0.987) | 6.39% | 0.830 (0.809-0.850) |
|  |  |  | Consensus | 8.42% | 0.904 (0.892-0.916) | 0.77% | 0.524 (0.379-0.670) |
|  |  |  | MetaSVM | 4.27% | 0.945 (0.934-0.957) | 4.92% | 0.811 (0.786-0.837) |
|  |  |  | MetaLR | 1.78% | 0.900 (0.874-0.925) | 7.41% | 0.871 (0.855-0.887) |
| TNNT2 | 1.54% | 0.912 (0.889-0.935) | HCM cluster | 1.23% | 0.958 (0.941-0.974) | 0.46% | 0.787 (0.699-0.874) |
|  |  |  | Consensus | 1.20% | 0.909 (0.880-0.937) | 0.49% | 0.832 (0.730-0.934) |
|  |  |  | MetaSVM | 1.11% | 0.894 (0.861-0.927) | 0.58% | 0.905 (0.848-0.961) |
|  |  |  | MetaLR | 1.11% | 0.889 (0.856-0.923) | 0.58% | 0.921 (0.872-0.971) |
| TNNI3 | 1.95% | 0.930 (0.912-0.948) | HCM cluster | 1.92% | 0.974 (0.963-0.984) | 0.18% (p=0.0918) | 0.457 (0.140-0.774) |
|  |  |  | Consensus | 1.93% | 0.957 (0.943-0.970) | 0.17% (p=0.0383) | 0.566 (0.280-0.852) |
|  |  |  | MetaSVM | 1.77% | 0.939 (0.921-0.957) | 0.33% | 0.873 (0.803-0.944) |
|  |  |  | MetaLR | 1.87% | 0.932 (0.913-0.951) | 0.23% | 0.903 (0.833-0.973) |
| MYL3 | 0.70% | 0.833 (0.772-0.895) | HCM cluster | 0.55% | 0.925 (0.886-0.965) | 0.29% (p=0.0021) | 0.655 (0.455-0.856) |
|  |  |  | Consensus | 0.79% | 0.869 (0.817-0.921) | 0.05% (p=0.6503) | 0.310 (0-1) |
|  |  |  | MetaSVM | 0.50% | 0.840 (0.763-0.917) | 0.34% | 0.833 (0.735-0.930) |
|  |  |  | MetaLR | 0.53% | 0.809 (0.722-0.897) | 0.31% | 0.883 (0.809-0.958) |
| CSRP3 | 0.41% | 0.683 (0.563-0.803) | HCM cluster | 0.43% | 0.882 (0.821-0.943) | 0.16% (p=0.5533) | 0.158 (0-0.724) |
|  |  |  | Consensus | 0.58% | 0.735 (0.630-0.839) | 0.02% (p=1.0000) | - |
|  |  |  | MetaSVM | 0.53% | 0.779 (0.687-0.871) | 0.07% (p=1.0000) | - |
|  |  |  | MetaLR | 0.55% | 0.751 (0.651-0.852) | 0.05% (p=1.0000) | - |

The performance of variant clustering and functional prediction scores in distinguishing between pathogenic and benign variants was then compared. In contrast to the significant enrichment of pathogenic variants obtained by analysis of the regional distribution of variation, functional prediction consensus scores only marginally increased EFs, compared to whole-gene estimates (Table 2), highlighting the limitations of using such generic predictors. For other HCM genes, no clear clustering of variants in the case cohorts was observed across the protein sequence (Figure S1). Therefore only consensus functional prediction scores are currently available for variant prioritisation, but again these provide only a marginal increase in EF values for these genes (Table S4).

### Adapting ACMG/AMP guidelines to incorporate EF prior probabilities

For non-truncating variants, there are currently two rules in the ACMG/AMP guidelines that can incorporate information on the differing frequencies of particular variant classes between case and control cohorts and that can be activated by novel variants – PP2 (*missense in gene with a low rate of benign missense variants and pathogenic missense variants are common*) and PM1 (*mutational hot spot or well-studied functional domain without benign variation*). However, activating even the stronger of these rules (PM1) will not lift any novel or relatively uncharacterised variant beyond VUS without substantial segregation or functional characterisation, even if found in genes or regions that are completely intolerant of variation. Additionally, the rules are categorical (despite describing a quantitative class of evidence) and must be specified for each gene and disease, with no consensus yet on the circumstances in which these should be applied.

In order to apply a more quantitative approach to these rules, we propose an adaptation of the guidelines as shown in Figure 2. The EF enables a unified approach and provides an empirical estimate of the probability of pathogenicity for a variant in a given gene (or region of a gene) that allows rules to be applied at different strengths. The non-quantitative related rules PP2 and PM1 would be replaced with a single rule (PM1) with three (or more) evidence levels depending on pre-defined EF for the relevant variant class. For genes where clustering of variants has been observed, regional EFs, rather than EFs at the gene level, should be applied. This semi-quantitative approach is similar to the PP1 rule for segregation data that allows the rule to be progressed from supporting to moderate to strong with increasing evidence[23, 24]. As the EF is calculated for rare variants found in cases, PM1 would only be activated in combination with the PM2 rule defining rarity. Since PM1_strong (in conjunction with PM2) would enable a novel variant to be classified as likely pathogenic, we suggest an EF≥0.95 could activate this rule. This is equivalent to an OR of 20, broadly similar to that adopted in the Bayesian modelling of the ACMG/AMP guidelines by Tavtigian et al[25].

**Figure 2:**
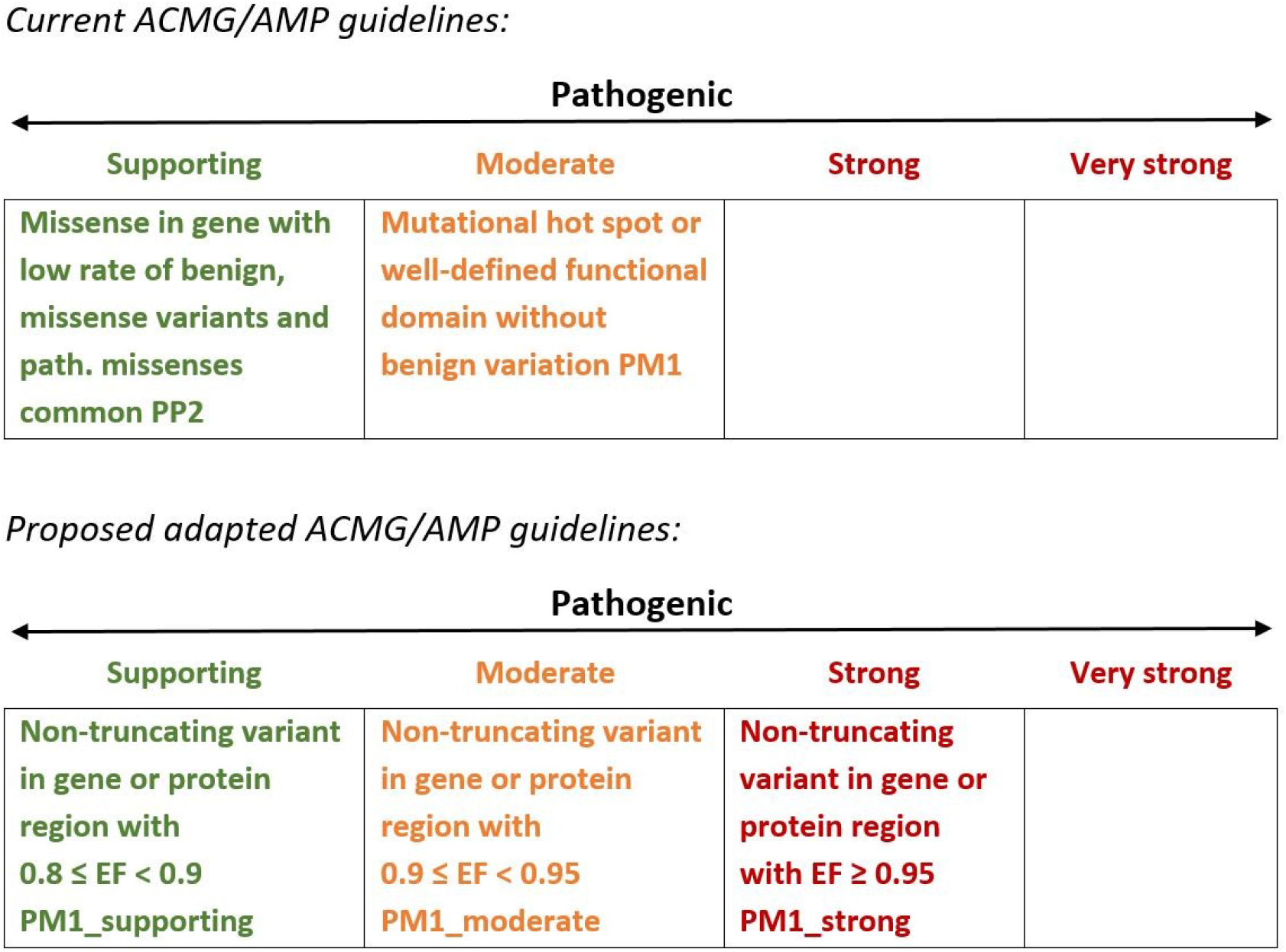
Proposed adaptation of ACMG/AMP guidelines relating to the relative frequencies of non-truncating variants in case cohorts and population controls.

Since each level of evidence in the hierarchical ACMG/AMP framework represents a doubling in weight, a Bayesian interpretation of the ACMG/AMP guidelines[25] requires that the odds should increase by a power of 2 as you move to a higher evidence tier. This yields corresponding EF/OR thresholds of 0.776/4.47 for the PM1_moderate rule and 0.527/2.11 for the PM1_supporting rule given an EF threshold of 0.95 for PM1_strong. However, we believe a more conservative application of these rules may be more appropriate in a real world setting, and therefore for this study we have defined PM1_moderate as an EF between 0.90 and 0.95 (minimum OR of 10) and PM1_supporting as an EF between 0.80 and 0.90 (minimum OR of 5). Future consensus-derived implementations of these rules may choose to incorporate the Bayesian model, although it should be noted that other recommendations for translating quantitative data into ACMG/AMP rules also do not account for exponentially scaled odds of pathogenicity[23, 24].

### An EF-calibrated tiered application of PM1 increases the yield of actionable variants in HCM

To evaluate how the EF-based modified ACMG/AMP guidelines could improve the yield of genetic testing in HCM, we determined the proportion of VUS in a diagnostic referral cohort that were found in genes or regions with an EF≥0.95, that might therefore trigger a PM1_strong rule (i.e. non-truncating variants throughout *TPM1* and in case-enriched clusters of *MYH7, MYBPC3, TNNI3* and *TNNT2*). In all, variants in 4.0% of cases could be upgraded to Likely Pathogenic by activating this strong evidence rule (Figure 3A). This represents an increase in yield of pathogenic and likely pathogenic variants in the eight sarcomeric genes from 28.8% to at least 32.8% (14% relative increase) in this cohort. It should be noted this is a conservative estimate, focusing only on PM1_strong, whereas variants activating PM1_moderate and PM1_supporting might also lead to a change in interpretation when combined with other lines of existing evidence.

**Figure 3:**
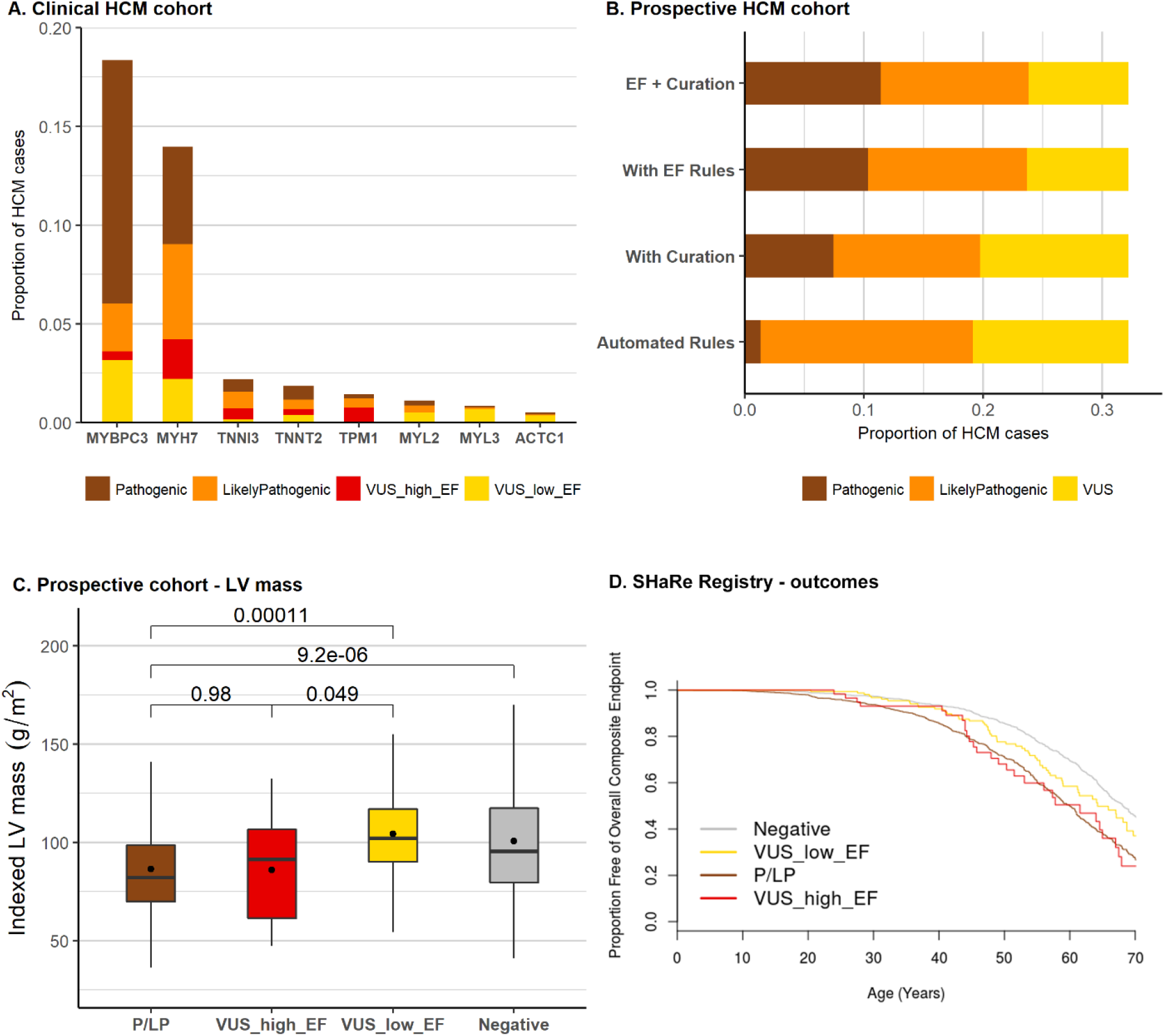
(A) Proportion of cases from the OMGL/LMM HCM cohorts with variants in 8 sarcomeric genes (only rare variants, ExAC filtering frequency < 4 x 10^−5^, are shown, excluding non-essential splice site variants). Coloured shading represents the clinical classification of the original diagnostic laboratory (OMGL and LMM), and, for variants originally classified as VUS, the proportion that could be reclassified as Likely Pathogenic based on occurrence within a gene or region with EF≥0.95. 89 variants in 123 cases for MYH7, 12 variants in 27 cases for MYBPC3, 18 variants in 34 cases for TNNI3, 15 variants in 18 cases for TNNT2 and 22 variants in 33 cases for TPM1 would be upgraded based on this analysis. (B) Proportion of cases in a prospective HCM cohort classified as actionable based on application of fixed and automatable ACMG/AMP rules, alongside the addition of manual curation of published evidence and the proposed EF-calibrated PM1 rules. 31 extra cases (4.5%) are upgraded with EF-based rules compared to just 4 (0.6%) with manual curation. (C) Comparison of indexed LV mass in cases with pathogenic variants, VUS in high EF (≥0.95) regions, and VUS in low EF regions (<0.95) in MYH7/MYBPC3 as well as genotype-negative cases, from the prospective HCM cohort. The clinical phenotype of individuals with VUS at locations anticipated to be pathogenic is indistinguishable from known Pathogenic/Likely Pathogenic variants, while individuals with VUS in other regions have a clinical phenotype more similar to individuals without a sarcomere variant. (D) Kaplan-Meier survival curve for the overall composite endpoint (including mortality, ventricular arrhythmia and heart failure composites) of the SHaRe cardiomyopathy registry stratified by genotype (HCM cases with pathogenic variants; VUS in high EF region (≥0.95); VUS in lower EF regions (<0.95); and genotype-negative cases).

Sarcomeric variants in a prospective cohort of 684 HCM cases[15] were also analysed. 19.1% of cases had actionable (pathogenic and likely pathogenic) variants with automatically applied rules (see Methods for details), with only 4 additional cases with VUS upgraded to actionable based on manual assessment of published evidence from family pedigrees. In contrast, VUS would be upgraded in 31 cases (of 82 with VUS) using the proposed PM1 modifications (4.5% of the cohort) in addition to automatically applied rules. In total, this corresponds to a 20.7% relative increase in actionable variants over current guidelines (Figure 3B). See Table S5 for details of the variants detected in this cohort.

### Independent validation of variants upgraded from VUS under this framework

The distinctive clinical characteristics of genotype-positive and genotype-negative HCM patients offer an opportunity to validate variant classifications in the absence of an independent gold-standard set of variants for benchmarking. If cases with variants that are upgraded from VUS to P/LP are more phenotypically similar to cases with known pathogenic variants, this offers further supportive evidence to validate the reclassification. We assessed mean indexed left ventricular (LV) mass and event-free survival as clinical variables that are associated with pathogenic sarcomere variants.

In the prospective HCM cohort, overall LV mass is significantly greater in genotype-negative cases (101.0±31.8g/m^2^) compared to genotype-positive cases (88.7±31.1g/m^2^), despite the fact that patients with pathogenic sarcomeric variants tend to have greater maximum LV wall thickness. Cases with variants upgraded from VUS were similar to genotype-positive (86.0±28.1 g/m^2^, p=0.98), with both significantly different from genotype-negative and cases with VUS that are not upgraded (104.3±24.7g/m^2^) (Figure 3C). Genotype-positive cases have significantly worse outcomes than genotype-negative cases, as demonstrated most comprehensively by data from the SHaRe registry (in press at Circulation). In this dataset, cases with VUS display intermediate outcomes, although more similar to genotype-positive (p=0.07) than genotype-negative (p<0.001). Sub-classifying these by EF, cases with VUS with an EF≥0.95 had similar outcomes to genotype-positive cases (p=0.9) and were significantly different to genotype-negative cases (p=0.001) (Figure 3D). In contrast, cases with VUS with an EF<0.95 displayed cumulative outcomes intermediate between genotype-positive (p=0.03) and genotype-negative (p=0.03) cases, consistent with the expectation that these cases will include a mix of both pathogenic and rare benign variants.

## DISCUSSION

The accurate and comprehensive interpretation of rare variants underlying Mendelian disease remains one of the principle challenges facing genetics and one of the key obstacles to fulfilling the potential of genomics in clinical practice. Current guidelines are conservative and prioritise minimising false positive results, given the potentially serious adverse consequences of predictive testing based on erroneously classified variants. However, this comes at the cost of sensitivity and denies many individuals the benefits of a molecular diagnosis. In HCM, case-control comparisons have highlighted that the majority of sarcomeric gene variants reported as VUS in leading clinical labs are in fact pathogenic variants, particularly for population groups that have not been extensively studied, highlighting the need for improved stratification of these variants. For applications of genetic testing other than diagnosis and predictive testing, such selection of specific therapies, a different balance between sensitivity and specificity may be required, and variants may be actionable with a lower burden of proof of causality. It is also recognised that VUS, though not clinically actionable, can create uncertainty and confusion for recipients of genetic testing, with patients often over-interpreting their effect[26]. New methods for more comprehensive identification of disease-causing variants, while maintaining the stringency of clinical guidelines, are urgently required.

In this study, we have demonstrated that using large case and population cohorts, and applying strict population frequency thresholds for variants of interest, we can identify genes and gene regions in which variants of specific classes have high likelihoods of pathogenicity. The probability of pathogenicity can also be empirically estimated, providing a quantitative measure of interpretative confidence. We demonstrate how the ACMG/AMP framework could be adjusted to incorporate this information (where suitable case series exist) and enable a more quantitative and transparent assessment of this evidence class. Crucially, this new framework allows variants that are novel or otherwise not yet well-characterised, but which belong to variant classes with very high prior probabilities of pathogenicity, to be classified as actionable. Under existing rules, such variants are condemned to remain as VUS unless the family structure permits well powered segregation analysis, or there are resources for functional characterisation.

As variant-specific evidence such as co-segregation data has typically been required to classify missense or non-truncating variants as disease-causing, we recognise that the novel approach to variant classification described here may require further piloting and replication before adoption a clinical setting. However, we believe this method is consistent with the stringent approach to variant classification of current guidelines. While the ACMG/AMP guidelines define likely pathogenic as a *“greater than 90% certainty of a variant being disease-causing”*[1], a 95% threshold is arguably more in line with standard clinical practice and therefore we have proposed an EF cut-off of 0.95 to define strong evidence for this rule. We consider a 95% probability of pathogenicity to be a reasonable level of evidence for a “likely pathogenic” classification, and one that provides an effective balance between sensitivity and specificity in genetic testing. It is also important to recognise that there is an inherent uncertainty associated with all variant interpretation, particularly for those classified as likely pathogenic. The confidence of both clinicians and patients in the results of genetic testing could be improved by more effective reporting of the evidence for pathogenicity in genetic reports, including the EF for relatively uncharacterised variants, and more transparency about the level of certainty associated with any classification.

Importantly, the approach to variant classification described here is compatible with the existing framework of the ACMG/AMP guidelines that have been widely adopted in clinical genetics laboratories. The translation of EF values into semi-quantitative PM1 rules, with a twofold increase in ORs required to progress between evidence classes, is similar to that adopted for another quantitative data type - co-segregation with disease in affected family members. Recent studies have sought to translate segregation data into supporting, moderate or strong PP1 rules based on the number of meioses of the variant that are informative for co-segregation[23, 24]. The rule adaptations proposed here also address the discrepancy between the rules for truncating and non-truncating variants in the current guidelines. Truncating variants in genes where loss of function is a known mechanism for the disease in question will achieve a classification of at least likely pathogenic, courtesy of the (very strong) PVS1 rule, assuming a number of criteria are met[27]. While the weight of this rule partly derives from the fact that a non-functional protein is likely to be produced by the truncating variant (albeit with the caveats described by Richards et al[1]), it also reflects the rarity of such variants in the population and consequently the high odds of a variant detected in a patient being pathogenic (as seen with *MYBPC3* truncating variants in this study, with an EF>0.99 and an OR of 115). Non-truncating variant classes that are similarly highly enriched in case cohorts should also have this evidence more appropriately weighted when evaluating variants.

Our findings highlight the necessity of applying gene and disease-specific expertise to both variant classification and the customisation of ACMG/AMP guidelines[6]. As we have shown, variant characteristics that are specific to the genes and disease in question, such as clustering of case variants in specific protein domains, are more powerful discriminators than generic techniques designed to be applied genome wide, such as the widely used missense functional prediction algorithms. This has also been recently demonstrated by an analysis of variation in the *RYR2* gene in catecholaminergic polymorphic ventricular tachycardia[28]. Interestingly, the estimated 14-20% increased yield of actionable variants in sarcomeric genes described here is likely to have a greater impact on HCM genetic testing than all of the efforts over the last 10-20 years to identify novel, non-sarcomeric genetic causes in this condition[15] that have explained very few additional cases. This highlights how efforts and resources to improve variant interpretation and the yield of genetic testing can be inefficiently allocated. While discovering valid novel genes may advance our understanding of disease and identify new therapeutic targets, an over-emphasis on discovering “novel” causes of diseases (from both researchers and journal editors/reviewers) may have less translational impact than efforts to improve our understanding of variation in known disease genes.

The publication and sharing of genetic data, as well as evidence about variant consequences in resources like ClinVar, is crucial for expanding our ability to interpret the results of clinical genetic testing of Mendelian disease[29]. This study also underscores the importance of clinical laboratories and research groups publishing and sharing genetic data with allele frequencies across cases cohorts as well as recording observations in individual patients – a large proportion of the HCM data in this study was published previously by the LMM[12] and OMGL[3] clinical laboratories. This will be even more critical for extending this approach to rarer and less well characterised genetic diseases than cardiomyopathies. However, the analysis of the prospective HCM cohort in this study has also exposed the limitations of relying on variant-specific evidence such as segregation data for the interpretation of variants. Published segregation data was mostly restricted to those variants that are already enriched in HCM cases (and therefore can be used to increase confidence in the variant classification by upgrading from likely pathogenic to pathogenic) rather than enabling rarer variants to be progressed from VUS to actionable, highlighting the necessity of novel approaches to increase the sensitivity of genetic testing, such as those described in this study.

### Comparisons with other methods that assess region pathogenicity

An alternative approach to identify functionally important genic regions seeks those that are depleted in (missense) variation in a reference population[22], in contrast to the analysis presented here that seeks a regional enrichment of variation in cases. Here depletion indicates negative selection of variation, implying that variation is not tolerated. Sub-genic regions of constraint were identified in only three of the HCM genes analysed in this study (*MYH7, FLNC, TNNC1*). There is partial overlap of the regions identified in this study (Figure 1), e.g. a region of high constraint in *MYH7* from residues 1-916 broadly corresponds to our HCM cluster (residues 167-931). Whatever the method for identifying a region of interest, empirical comparison of cases and controls provides a direct assessment of the strength of association with a specific disease, enabling us to directly estimate the likelihood of pathogenicity for variants in specific regions, as well as detecting pathogenic clusters in other genes for which no regional constraint data exists.

The EF (and OR) can of course be applied to calibrate an appropriate PM1 rule strength irrespective of the method by which the region is initially highlighted as potentially important. For example, two recent studies explored structure-function models in β-cardiac myosin (*MYH7*) to identify residues that are key to protein function (and therefore intolerant of variation), with variants affecting these residues enriched in case over population reference cohorts. Homburger et al modelled β-cardiac myosin before and after the myosin power stroke and identified the converter kinetic domain and myosin mesa surface area as regions enriched in disease-associated variants using a spatial scan statistic[20]. Alamo et al defined sites of inter- and intramolecular interaction between pairs of myosin heads (the interacting-heads motif - IHM), noting that variants in HCM cases disproportionately alter IHM residues[21]. The *MYH7* residues identified by these studies largely overlap with the HCM cluster we have identified by one-dimensional clustering (Figure 1). Particular groups of residues detected by these analyses are highly enriched in disease-associated variants (yielding higher EFs than our cluster), with 7 IHM groups yielding an EF>0.99 and accounting for 44% of variants found in HCM cases[21]. EF-based variant analysis thus requires a balance between specificity and sensitivity, or a tiered approach with different confidence levels for pathogenicity.

### Issues and limitations of this approach to variant classification

The calculation of EFs for particular variant classes is dependent on a number of factors. As we have previously shown, it is critical to adopt stringent, disease-specific frequency thresholds when assessing putative pathogenic variants[14]. Additionally the choice of case and control/population cohorts will influence how EFs are generated. For cases, the use of clinical referral cohorts (the diagnostic case series from different centres in Europe and North America that we have employed in this study should be reasonably representative of real-world referral patterns)) will produce more conservative EF values than highly selected case series, but we believe these more cautious, referral EFs are relevant to a clinical genetic setting. Nonetheless these may change as referral patterns change (e.g. with increasing test availability). Although the use of population reference data without well-defined phenotypes has limitations, and is not optimally matched technically (e.g. differences in sequencing coverage, as previously discussed[3]), we believe the advantages (population size and ethnic diversity allowing more accurate calculation of rare variant frequencies) outweigh the disadvantages. It is also crucially important to note that the evidence described here should only be applied to assessing variants from patients with the disease in question, and not from incidental or secondary findings in healthy individuals or those being sequenced in the context of other conditions, as EFs correspond to the probability of pathogenicity *given that the variant is identified in an individual with disease*.

Although we have identified highly pathogenic variant classes in a number of HCM genes, for others it is more challenging to effectively differentiate between benign and pathogenic variation. In particular, although we detected some small case-enriched clusters of non-truncating variants in *MYBPC3*, these will correspond to only a small proportion of such variants that are responsible for HCM in up to 8% of cases. For such genes, further research and larger datasets are needed to identify the protein regions and specific residues at which variation is most likely to cause disease. This could include analysis of protein structure, as demonstrated in the *MYH7* studies described above, or the development of computational prediction techniques that are specific to (and validated in) key disease genes, given the limitations of the generic and consensus scores that we have observed in this study. Genes like *MYBPC3*, i.e. those with a high diagnostic yield but poor signal-noise ratio that impedes the statistical prediction of pathogenicity, could also be prioritised for high-throughput functional classification studies[30].

### Conclusion

In conclusion, we have demonstrated that by combining large case and control datasets, stringent population frequency thresholds and the detection of pathogenic clusters in key disease genes, we can empirically estimate the likelihood that rare variants in specific genes or regions are pathogenic and can identify variant classes with a high prior probability of pathogenicity. Using this evidence to calibrate the appropriate weighting for rules within the ACMG/AMP framework, we believe the yield of genetic testing in diseases like HCM can be significantly increased, with less dependence on the prior characterisation of variants to define pathogenicity, while retaining a robust statistical framework. This may help to reduce the ethnicity bias associated with obtaining a positive result, enabling a more equitable application of genetic testing. This study also reinforces the concept that disease and gene-specific approaches are critical for accurate and comprehensive variant analysis. Finally, this quantitative approach moves us towards more transparent probabilistic variant classification for both for Mendelian disease genetics and precision medicine.

## METHODS

### Calculation of etiological fraction for significantly enriched variant classes

The etiological fraction (EF) estimates the proportion of risk that can be attributed to a specific exposure, in a population with disease who have been exposed to a risk factor[3]. In the context of Mendelian disease, exposure refers to a rare protein-altering variant in a particular gene, and the EF estimates the proportion of cases with a rare variant in whom that variant is disease-causing. The EF is derived from the attributable risk percent (ARP) among exposed, i.e. expressing the risk as a proportion rather than a percentage, and derived from the odds ratio (OR) as described below, where the OR provides an accurate estimate of the relative risk (RR) – the ratio of risk among exposed to risk among unexposed[31]. The odds ratio (OR) is calculated by (Altman, 1991)[32]:

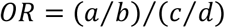

where a = disease cases with a variant, b = controls/reference population with a variant, c = disease cases without a variant, d = controls/reference population without a variant. The 95% confidence intervals (CI) for OR values are calculated by:

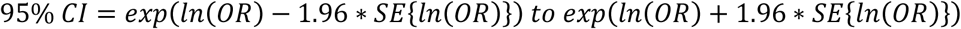

where the standard error of the log OR was given by:

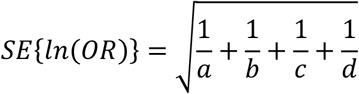

The EF is derived from the OR:

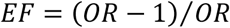

95% CIs for EF values are calculated as described by Hildebrandt et al[33].

EF and OR values were calculated for both truncating (frameshift, nonsense, splice donor site, splice acceptor site) and non-truncating (missense, small in-frame insertions/deletions) variants in HCM genes where a significant excess of rare variants in cases over the ExAC reference population was observed[15]. For the eight core sarcomeric genes (*MYBPC3, MYH7, TNNT2, TNNI3, TPM1, MYL2, MYL3, ACTC1*), the case cohorts were derived from published data from the Oxford Molecular Genetics Laboratory (OMGL) and the Laboratory of Molecular Medicine (LMM), Partners Healthcare, comprising between 4185 and 6179 unrelated HCM probands[3, 12]. For the minor HCM genes (*CSRP3, FHL1, PLN, TNNC1*), combined cohorts from OMGL and LMM, a prospective research cohort from our laboratory and published cohorts were used as previously described[15], comprising between 2061 and 5440 unrelated HCM probands. For *FLNC*, a recently published cohort of 448 HCM patients was used[19]. All rare variants were included for these calculations, regardless of the clinical classification of the variants.

For all genes, ExAC was used as the reference population database for background variation as previously described[3]. To account for variable coverage of the exome sequencing in ExAC, the sample total for each gene was adjusted by calculating the mean number of called genotypes for each variant. Rare variants were defined as those with a filtering allele frequency in ExAC below the maximum credible allele frequency for HCM[14], defined as 4x10^−5^ (prevalence = 1 in 500, allelic heterogeneity = 0.02, penetrance = 0.5, monoallelic inheritance, as calculated at http://cardiodb.org/allelefrequencyapp/).

### EFs as a means of quantifying performance of variant classifiers

The EF is dependent on the relative frequencies of variants in cases and population controls. While applying strict thresholds for rarity will focus on variants more likely to be disease-causing, thereby increasing the EF, this is usually not sufficient to adequately distinguish between benign and pathogenic variation for non-truncating variants. Therefore additional methods are required to discriminate between causative and background variants. A perfect discriminator of pathogenic and benign variants will identify the proportion of causative variants that is equal to the case excess and yield an EF of 1.0, with the proportion of benign variants equal to the population reference frequency of ExAC (and an EF of 0) – see hypothetical example in Figure 4. In practice, it is unlikely that full discrimination will be achieved but this EF-based approach allows us to evaluate methods that aim to differentiate between pathogenic and benign variants. In this study we compare the widely used and generic missense functional prediction scores with gene and disease-specific variant clustering. This EF-based approach also offers the advantage of not requiring predefined lists of irrefutable pathogenic and benign variants, which can be limited when performing analyses on specific genes.

**Figure 4:**
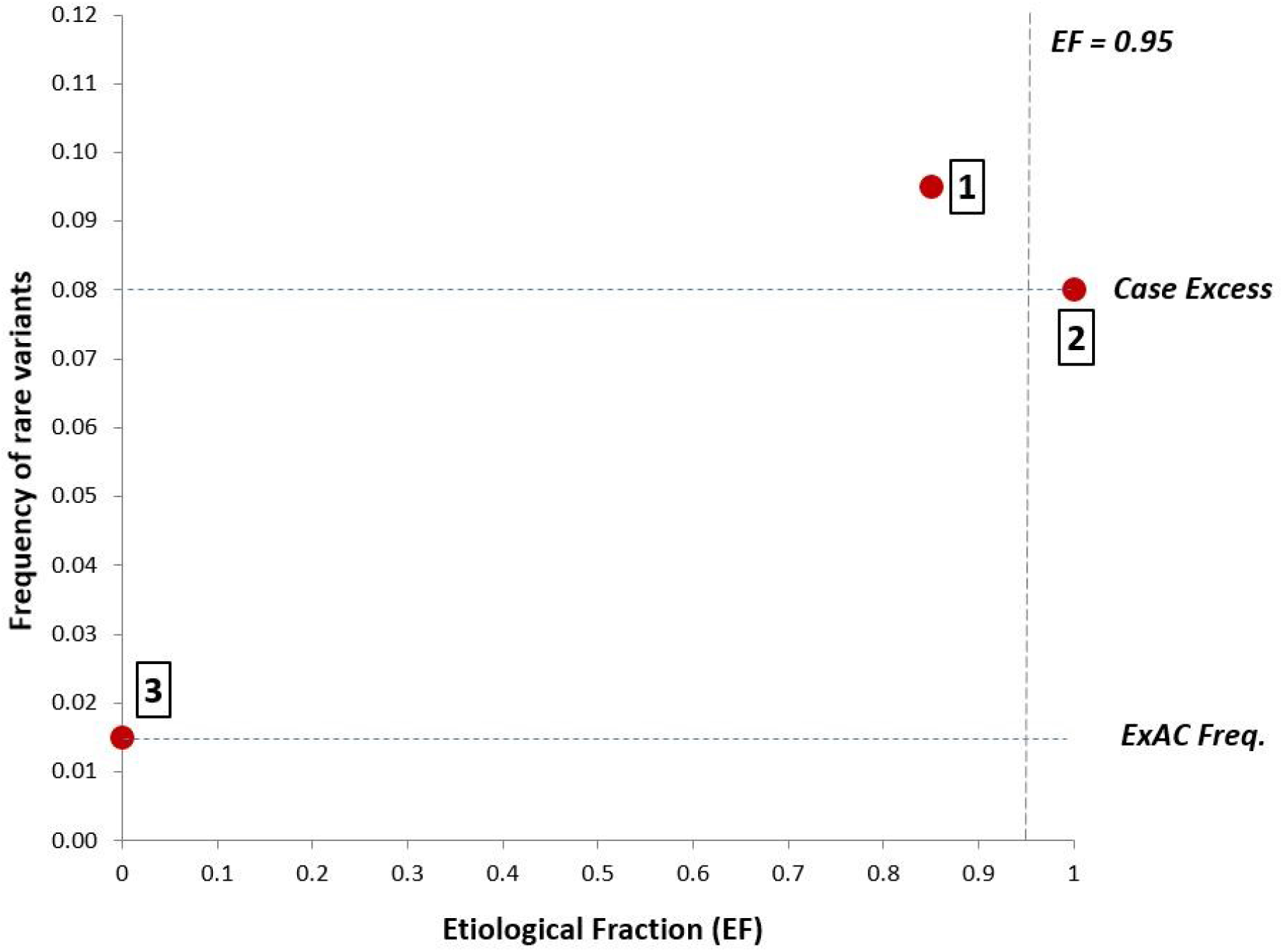
Illustration of how EFs can be used to evaluate methods for distinguishing pathogenic from benign variants (for a hypothetical gene). The overall EF of 0.85 [1] is based on a case frequency of 9.5% and a reference frequency of 1.5%. The aim of variant classification methods is to fully distinguish between pathogenic variants (producing an EF of 1.0 with frequency equal to case excess [2]) and benign variants (producing an EF of 0 with frequency equal to population reference, here ExAC [3]). We propose that an EF of 0.95 would be required to indicate a likely pathogenic variant.

### Assessing performance of missense functional prediction scores in HCM genes

Functional prediction scores from the dbNSFP database[17] (version 3.2) were downloaded for all missense variants in the 13 HCM genes. Eight scores that provide binary predictions, i.e. damaging vs benign/neutral were assessed – fathmm-MKL coding, FATHMM, LRT, Mutation assessor, MutationTaster, Polyphen2-HDIV, PROVEAN and SIFT, as well as the CADD algorithm (damaging variants were defined with a CADD phred score ≥ 15). A consensus prediction between the 9 scores was defined as being damaging if greater than 50% of the scores predicted a damaging effect. Additionally two consensus algorithms, MetaLR and MetaSVM[16], were also evaluated. The proportion of available predictions for each score for all potential missense variants in each gene was calculated to identify algorithms that do not provide comprehensive predictions for specific genes.

To test the effectiveness of these prediction scores for individual HCM genes, missense variants of known consequence (pathogenic and benign missense) identified. Pathogenic variants were defined by rarity in ExAC as described above and:

1. classified as pathogenic (P) or likely pathogenic (LP) in HCM patients by two or more clinical laboratories (OMGL, LMM and ClinVar submitters).
2. classified as P/LP by one clinical laboratory with no conflicting classifications (VUS or benign) by other laboratories.
3. significantly enriched in the OMGL/LMM cohorts compared to ExAC (Fisher’s exact test).

Benign variants were defined as:

1. presence in more than one individual in ExAC and not associated with any disease in ClinVar (P/LP/VUS) or HGMD
2. associated with disease in ClinVar (though not P or LP) or HGMD but at a frequency >0.001 in ExAC.

The sensitivity (true positive rate) and specificity (true negative rate) was calculated for the 9 functional prediction scores and 3 consensus scores for each of the 8 core sarcomeric genes (there were insufficient known pathogenic variants for the minor genes). As an alternative method for assessing these predictors, EFs were calculated for deleterious variants using the case and ExAC cohorts described above.

### Clustering algorithm to detect regional enrichment of variants

Protein regions enriched for rare variants were identified using a bespoke unsupervised clustering algorithm developed within this project. The algorithm is based on a sliding window scanning the protein sequences from their N-terminal to C-terminal residues, with a binomial test used to detect whether there is significant variation enrichment within the tested window compared with the rest of the protein.

The results of this first step are influenced by the size of the sliding window, with a spectrum ranging from small windows enabling detection of smaller, highly enriched variation hotspots but prone towards overfitting (in the most extreme case each residue with multiple variant alleles is considered a cluster), to large windows enabling detection of more extended enriched regions such as large protein domains but at the risk of too low a resolution (in the most extreme case, a unique cluster starting at the first variant residue and ending at the last). In terms of model performance, the former situation is characterized by specificity=1 (no variant-free residues are within clusters) and sensitivity close to 0 (the vast majority of variant residues are excluded from clusters), whereas the latter results in the opposite situation (many variant-free residues are included in the unique cluster [specificity close to 0] but also all variant amino-acids are [sensitivity=1]). For this reason, the algorithm automatically selects the optimal window size for each protein by searching for one minimizing the difference between sensitivity and specificity (in this case the mean difference between cases and controls for each gene). Of note, the sparseness of the data (resulting in a strong imbalance between positive data points [variant residues] and negative data points [variant-free residues]) make all classic model performance measures (e.g. accuracy, AUC, PPV etc.) biased towards results obtained with smaller window sizes.

To look for the optimal window size, the algorithm starts by testing 19 different sizes ranging from 5% of the protein to 95%. Subsequently, the algorithm picks the best one (if any) and tests 18 sizes around it at a 10-fold finer resolution (e.g. if the initial best window size is 10%, the next iteration will be on windows between 5.5% and 14.5%). This iterative process is repeated until a performance plateau is reached (i.e. none of the 18 new window sizes decreases the difference between sensitivity and specificity by more than 0.001 compared with the previous iteration). Once the optimal window size is detected, multiple testing correction is applied to each definitive window significantly enriched for variation, on the basis of the average number of times each protein residue has been tested (which depends on the number of iterations made, and on the size of the tested windows). Whenever a significant enrichment is detected within a window, its coordinates (start/end) are stored until the whole protein is scanned and, subsequently, merged with any other significantly enriched window to obtain a first “raw” set of variation-rich clusters.

After this first step, the algorithm performs a “boundary trimming” procedure at both ends of each cluster. This step controls for potential inclusion of variant-free (or non-enriched) distal cluster tails that may have been included within a significantly enriched window due to variants occurring more proximally. The algorithm performs the same procedure at both the N- and the C-terminal cluster boundaries, starting with a single-residue window including only the most external amino-acid, and iteratively extending it as far as the cluster median residue. Before each extension, the binomial test is used to check if there is a significant depletion of variants compared to the rest of the cluster. The algorithm stores each test’s p-value and tested region coordinates, and eventually trims the cluster by removing the most (if any) significantly variation-depleted tail, to obtain a final, refined set of clusters. One last binomial test is performed on the refined clusters to measure the significance of their rare variant enrichment.

### Distinguishing pathogenic from benign variants using clustering in case and control cohorts

EFs were calculated based on these clusters and compared to those produced by a consensus of missense functional prediction scores from the dbNSFP database[17] (MetaLR, MetaSVM and a consensus of 9 individual predictors as described above). These consensus scores were also evaluated in genes where no clustering of case variants was observed.

### Using EFs to increase the yield of putatively pathogenic variants in HCM cohorts

Sarcomeric gene rare variants in the OMGL/LMM clinical cohort[3] were re-assessed based on the analysis described above. The proportion of patients with variants that would be upgraded to Likely Pathogenic based on the revised ACMG/AMP guidelines was calculated, i.e. those previously classified as VUS but in a variant class with an EF≥0.95 for missense variants or EF≥0.90 for inframe indels (as inframe indels will also activate the PM4 rule regarding variants that change protein length and therefore only the moderate PM1 rule would be required for a likely pathogenic classification).

### Analysis of prospective HCM cohort

The effect of the new EF-based ACMG/AMP rules on the yield of actionable variants was assessed on a prospective cohort of 684 HCM patients recruited at the Royal Brompton & Harefield Hospitals NHS Foundation Trust, London UK[15]. The ACMG/AMP rules described below were used to classify variants from the valid HCM genes defined in this study, with rule implementation as described in the CardioClassifier resource[6]. The following rules could be activated by automated script:

- PM2 – filtering allele frequency in ExAC < 4 x 10^−5^. This rule must be activated to denote a causative variant for this analysis.
- PVS1 – truncating variants in *MYBPC3, TNNT2, TNNI3, CSRP3, FHL1, PLN* (genes statistically enriched in HCM cohorts versus ExAC).
- PS4 – individual variant statistically enriched in cases over controls, based on LMM/OMGL cohort versus ExAC with the rule activated if the case count was >2 and the Fisher’s exact test p-value < 1.79x10^−6^ (Bonferroni correction).
- PM4 – protein length changing variant, i.e. an inframe indel or stop lost variant.
- PP3 – missense variant with multiple lines of computational evidence suggesting a deleterious effect, i.e. of the 8 predictors assessed (SIFT, PolyPhen2 var, LRT, Mutation Taster, Mutation Assessor, FATHMM, CADD and Grantham scores), only 1 predicts benign and <3 have unknown classifications or if ≥3 have unknown classifications, all others predict damaging.
- PM5/PS1 – novel missense change at an amino acid where a different missense variant is pathogenic (PM5) or novel missense variant with same amino acid change as an established pathogenic variant (PS1). Pathogenicity here is defined as a pathogenic classification in ClinVar by multiple submitters with no conflicting evidence.

Rare variants (i.e. with rule PM2 activated) were then manually assessed for human genetic evidence in ClinVar entries and published reports using the following rules:

- PP1 – co-segregation with disease. This rule was defined as supporting for ≥3 observed meioses, moderate for ≥5 meioses and strong for ≥7 meioses.
- PS2/PM6 – de novo inheritance (with/without confirmed paternity and maternity).
- The PS3 rule relating to effects in functional studies was not applied due to the lack of standardisation and validation in functional assays for HCM variants.

The number of patients with variants that still remained as VUS, i.e. unactionable according to current guidelines, but that would be upgraded to at least Likely Pathogenic based on the revised ACMG/AMP guidelines was calculated as described for the clinical HCM cohort, i.e. those in a variant class with an EF≥0.95 for missense variants (activating PM1_strong) or EF≥0.90 for inframe indels (activating PM1_moderate).

### Genotype-phenotype analyses to validate variant pathogenicity

The clinical characteristics of two HCM cohorts were used to support the pathogenicity of variants upgraded on the basis of an EF≥0.95. For the prospective HCM cohort, left ventricular (LV) mass values indexed to body surface area were derived from cardiac magnetic resonance imaging and compared between cases with pathogenic or likely pathogenic variants (current ACMG/AMP guidelines), VUS upgraded to likely pathogenic with EF rules, other VUS and genotype-negative cases (only variants in thick filament genes *MYH7* and *MYBPC3* were analysed due to the distinctive patterns of LV hypertrophy observed in cases with variants in thin filament genes[34]).

Outcome data was assessed using the Sarcomeric Human Cardiomyopathy Registry (SHaRe), a multicentre international repository that aggregates clinical and genetic data from patients with cardiomyopathies including HCM. A total of 2694 HCM patients with both right-censored outcome data and known sarcomeric genotype were analysed - 1254 patients with at least one pathogenic or likely pathogenic variant in any of the 8 sarcomeric genes; 1199 patients with no sarcomeric variants; and 241 patients with VUS in any of the sarcomeric genes. Of the 241 patients with VUS, 69 were reclassified as pathogenic as they had variants with an EF≥0.95. Survival curves were calculated by Kaplan-Meier analysis with log-rank test for the proportion of patients free of the overall composite outcome [in press at Circulation] for each of the four genotype groups.

---

**Figure S1:**
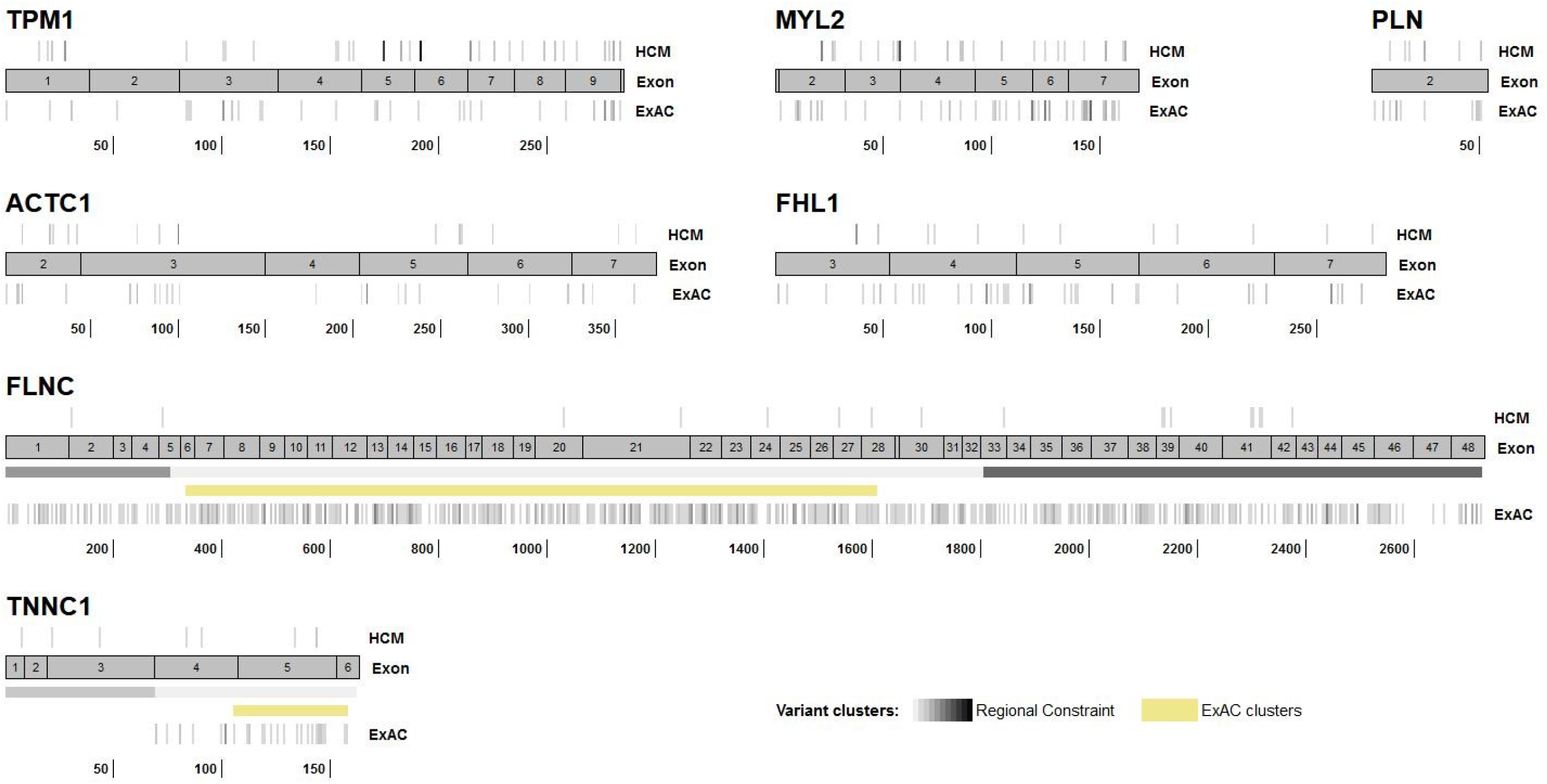
Distribution of rare, non-truncating variants in HCM cohorts and ExAC for validated genes without observed clustering of variants in cases (variant density increases with darker shades of grey). Regional constraint boundaries described by Samocha et al are also highlighted.

